# Cdk5 drives formation of heterogeneous pancreatic neuroendocrine tumors

**DOI:** 10.1101/2021.05.25.445594

**Authors:** Angela M. Carter, Nilesh Kumar, Brendon Herring, Chunfeng Tan, Racheal Guenter, Rahul Telange, Wayne Howse, Fabrice Viol, Chris Graham, Tyler R. McCaw, Hayden Bickerton, Frank Gillardon, Eugene A. Woltering, Deepti Dhall, John Totenhagen, Ronadip Banerjee, Elizabeth Kurian, Sushanth Reddy, Herbert Chen, Joerg Schrader, J. Bart Rose, M. Shahid Mukhtar, James A. Bibb

**Affiliations:** Department of Surgery, University of Alabama at Birmingham, Birmingham, AL 35233, USA; Department of Biology, University of Alabama at Birmingham, Birmingham, AL 35233, USA; Department of Psychiatry, University of Texas Southwestern Medical Center, Dallas, TX 75390, USA; Department of Internal Medicine, University Hospital Hamburg-Eppendorf, Hamburg, Germany; Department of Medicine, University of Alabama at Birmingham, Birmingham, AL 35233, USA; Boehringer Ingelheim Pharma GmbH & Co. KG, CNS Diseases Research, Birkendorferstrasse 65, 88397 Biberach an der Riss, Germany; Department of Surgery, Louisiana State University Health Sciences Center, New Orleans, LA 70112, USA; Department of Anatomic Pathology, University of Alabama at Birmingham, Birmingham, AL 35233, USA; Department of Radiology, University of Alabama at Birmingham, Birmingham, AL 35233, USA; Department of Pathology, University of Texas Southwestern Medical Center, Dallas, TX 75390, USA; O’Neal Comprehensive Cancer Center, Alabama at Birmingham, Birmingham, AL 35233, USA

**Keywords:** cyclin-dependent kinase, pancreatic neuroendocrine tumors, inducible mouse model, tumorigenesis

## Abstract

Pancreatic neuroendocrine tumors (PanNETs) are a heterogeneous population of neoplasms that arise from hormone-secreting islet cells of the pancreas and have increased markedly in incidence over the past four decades. Non-functional PanNETs, which occur more frequently than hormone-secreting tumors, are often not diagnosed until later stages of tumor development and have poorer prognoses. Development of successful therapeutics for PanNETs has been slow, partially due to a lack of diverse animal models for pre-clinical testing. Here, we report development of an inducible, conditional mouse model of PanNETs by using a bitransgenic system for regulated expression of the aberrant activator of Cdk5, p25, specifically in β–islet cells. This model produces a heterogeneous population of PanNETs that includes a subgroup of well-differentiated, non-functional tumors. The utility of this model is enhanced by ability to form tumor-derived allografts. Production of these tumors demonstrates the causative potential of aberrantly active Cdk5 for generation of PanNETs. Further, we show that human PanNETs express Cdk5 pathway components, are dependent on Cdk5 for growth, and share genetic and transcriptional overlap with the INS-p25OE model. This new model of PanNETs will facilitate molecular delineation of Cdk5-dependent PanNETs and the development of new targeted therapeutics.

## Introduction

Pancreatic neuroendocrine tumors (PanNETs) are a diverse group of neoplasms that originate from islet cells of the pancreas^1^. These tumors have the potential to secrete a range of bioactive hormones such as insulin, glucagon, and somatostatin. Tumors that secrete quantities of hormones that result in elevations in blood plasma levels are classified as functional^2^. Functional tumors produce hormonal syndromes commensurate with the hormone produced in excess^3^. Functional tumors are typically lower in grade and have good prognoses, possibly due to early detection as a result of the syndromes experienced by patients^4^. However, the majority of PanNETs are non-functional, which on average have comparatively worse prognoses^5^. Historically rare, the incidence of PanNETs in the United States increased 8-fold from 1973 to 2012^4^. Surgical resection provides excellent outcomes and long-term survival for patients with early stage primary tumors^6-8^. However, many PanNETs are metastatic at diagnosis and there are no curative therapies for advanced disease^9,10^.

Multiple molecular alterations have been implicated in the development of PanNETs. Mutations in the gene *MEN1* occur in approximately 40% of PanNET patients and changes in *DAXX/ATRX* are present in another 40%. Roughly 15% of patients possess changes that target the mTOR pathway, including mutations in *TSC2, PIK3CA*, or *PTEN*^11-13^. Unfortunately, thus far, no correlation has been observed between the presence of these mutations and patient response to specific pathway-targeted therapies in NET clinical trials^14^. Recently, cyclin-dependent kinase 5 (Cdk5) was implicated in the growth of several types of neuroendocrine tumors including PanNETs^15-17^. Interestingly, the presence of a set of downstream biomarkers of Cdk5 pathway activation was predictive of tumor growth inhibition in preclinical testing of a Cdk5-targeted therapy^17^.

Cdk5 is a non-canonical member of the Cdk family of proline-directed serine/threonine kinases^18^. Traditional family members, such as Cdk1, 2, 4, and 6, are important cell cycle regulators that are activated by cyclins and required for cell division^19^. Unlike these family members, Cdk5 is not activated by cyclins and is not required for normal cell division. Instead, Cdk5 is regulated through binding to cofactors p35 or p39^20,21^. The resulting protein complex plays a prominent role in several physiological processes in neuronal cells, such as proper migration for normal CNS development and function^22,23^. Interestingly, aberrant activation of Cdk5 has been implicated in several neurodegenerative diseases^21^. The pathological role of Cdk5 is facilitated through calpain cleavage of p35 to p25, a highly stable fragment that exhibits mislocalization in cells but retains the ability to bind and activate Cdk5^24,25^. Cdk5 pathway components are also expressed in neuroendocrine cells of pancreatic islets where they contribute to normal hormone secretion and β–cell survival^26-30^. New studies show that under conditions of aberrant activation in non-neuronal cells, Cdk5 can hijack signaling components traditionally involved in the cell cycle and successfully promote proliferation and/or migration^15,31-36^. Here, we show that Cdk5 and its activators are retained in islet cells that develop into PanNETs in humans and that aberrant activation of Cdk5 is involved in human PanNET cell growth. Furthermore, we show the potential for Cdk5 to drive development of PanNETs by demonstrating that expression of the aberrant activator, p25, in islets of mice, initiations tumor formation. Importantly, these PanNETs exhibit a heterogeneous phenotype that includes both functional and non-functional, well-differientated tumors.

## Results

To better understand the relevance of the Cdk5 pathway to human PanNETs, we performed immunostaining on distinct groups of grade 1 human tumors for Cdk5 pathway components. This revealed the presence of Cdk5 and its activators, p35 and/or p25 (p35/p25) (**Fig. 1A**) in both functional and non-functional tumors. To gain further insight into the prevalence of these signaling proteins in the PanNET patient population, we performed immunostaining on a PanNET tissue microarray (TMA) composed of 23 grade 1 tumors, 13 grade 2 tumors, 1 grade 3 tumor, and 5 different normal tissue controls. (**Fig. 1B-C** and Supplementary Table S1). Semi-quantitation revealed clear expression of Cdk5 and p35/p25 throughout these grades of tumors (**Fig. 1D-E**) and elevated expression relative to a normal placenta control sample.

**Figure 1.**
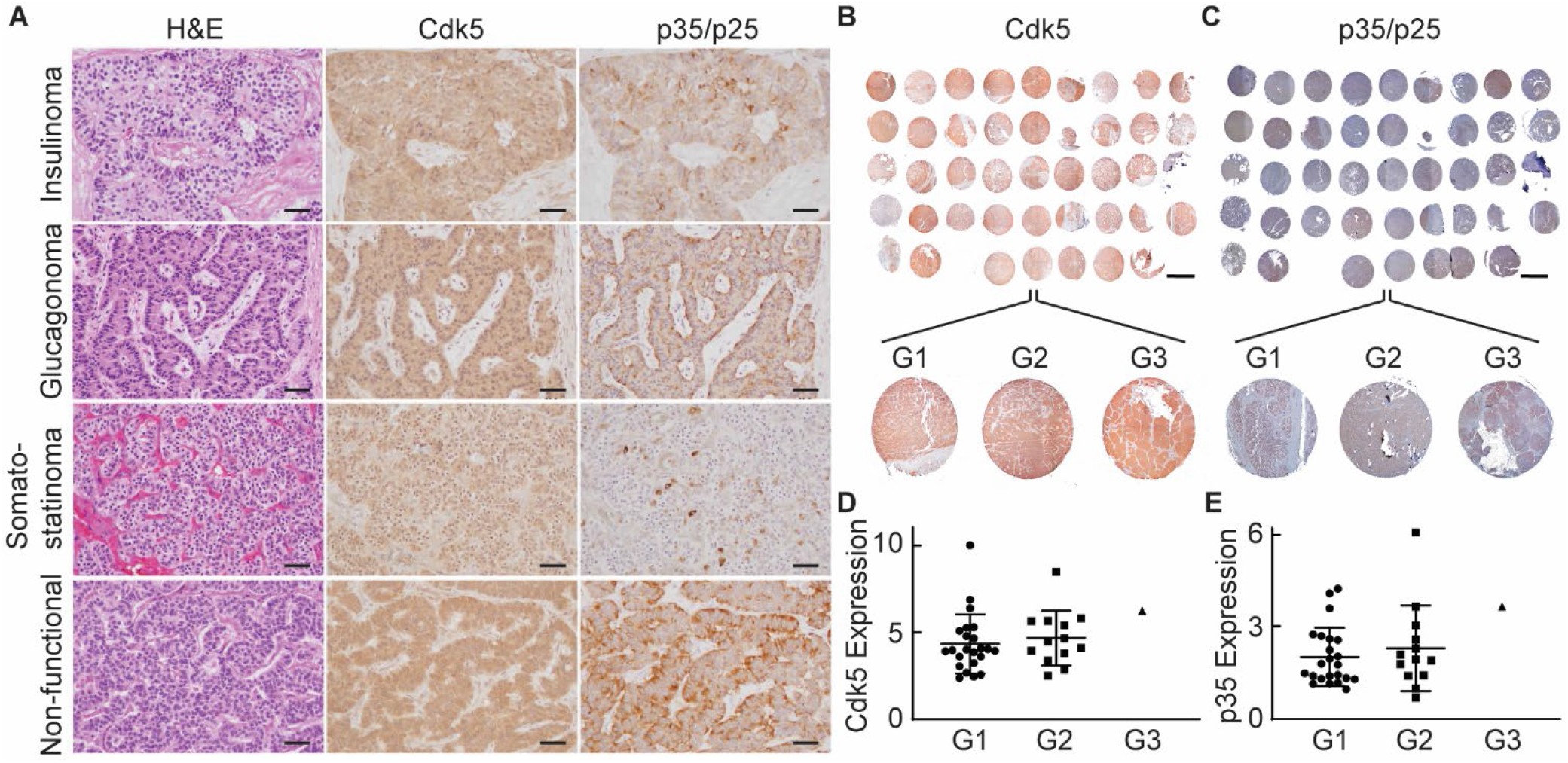
Cdk5 pathway components are present in human PanNETs. A. H&E stain and immunostains for Cdk5 and p35/p25 in G1 PanNETs. Scale bars = 50 μm. B-C. Immunostains for Cdk5 (B) and p35/25 (C) from human PanNET TMA. Scale bar = 2 mm. D-E. Semi-quantitation of Cdk5 expression from B (D) and p35/25 expression from C (E) normalized to expression levels of each in a normal placenta core; column 1 row 4 of MA. Map of TMA in Supplementary Table S1.

To determine if Cdk5 and it’s activators play a functional role in PanNETs, we next examined a set of human PanNET cell lines including the well-established BON and QGP lines, and two newly derived lines NT-18P and NT18-LM^37^. All cell lines expressed Cdk5 and its aberrant activator, p25 (**Fig. 2A**). We previously found that growth of the pancreatic carcinoid cell line, BON, was blocked by 4 different selective Cdk5 inhibitors and not by Cdk2 and Cdk4 specific inhibitors^17^. Here, we show that growth of all five PanNET cell lines tested is inhibited by the Cdk5-selective inhibitor, IndoA (**Fig. 2B**). These data indicate that Cdk5 dependence is a common feature shared by many PanNETs.

**Figure 2.**
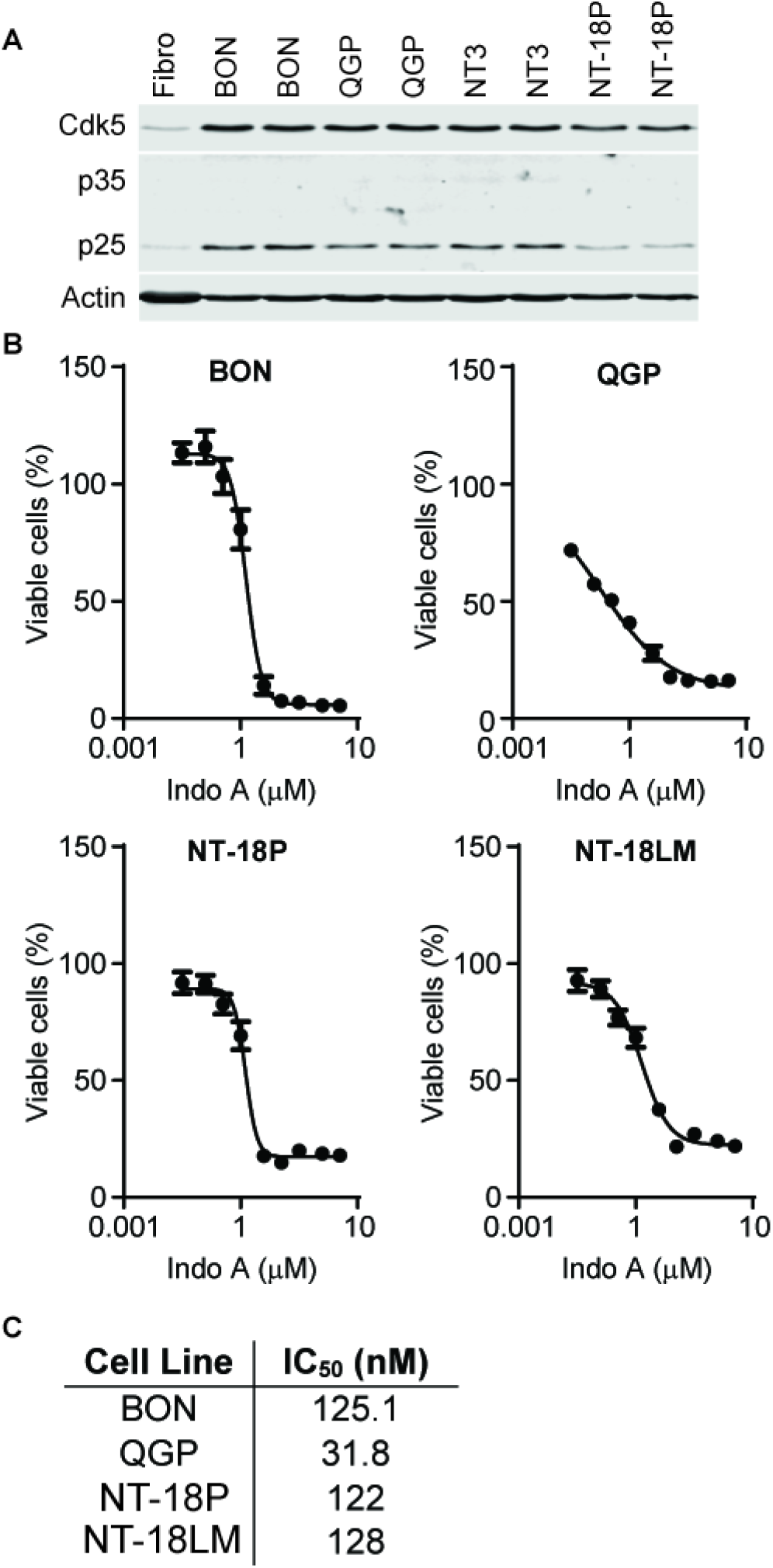
Human PanNET cells are dependent on Cdk5 for growth. A. Immunoblot of Cdk5 pathway components in fibroblasts and PanNET cells. B. PanNET cell lines were treated with increasing concentrations of Indo A and monitored for effects on cell viability. Error bars represent SEM. C. IC_50_ values obtained from viability assays in B.

To determine if Cdk5 has the potential to behave as a causative factor in PanNET tumorigenesis, we generated a bitransgenic mouse line in which expression of the aberrant Cdk5 activator, p25, can be induced in β-cells of the pancreas by addition of the small molecule doxycycline (dox) to drinking water. This was achieved by crossing the Ins2-rtTA mouse line^38^ that expresses the reverse tetracycline transactivator under the control of the insulin promoter with the tetOp-p25GFP line^39^ that expresses p25GFP under the control of the tetOp promoter (**Fig. 3A**) to produce bitransgenic offspring (INS-p25OE). As previously observed with some doxycycline (dox) inducible systems, a low level of transgene expression was observed in the absence of dox. However, administration of 1 g/L dox to INS-p25OE animals for 4-8 weeks further induced expression of the p25-GFP transgene in pancreatic islets (**Fig. 3B-C**). Formation of solid lesions in the pancreas were observed as early as 6 months post-induction of p25GFP expression (**Fig. 3D** and Supplementary Fig. S1). As confirmation that transgene expression does not occur ubiquitously throughout tissues of these animals, we examined samples of pancreatic masses along with liver and kidney tissues for p25GFP expression after 12 months of dox administration and found no evidence of p25GFP expression in non-pancreatic tissues (**Fig. 3E**).

**Figure 3.**
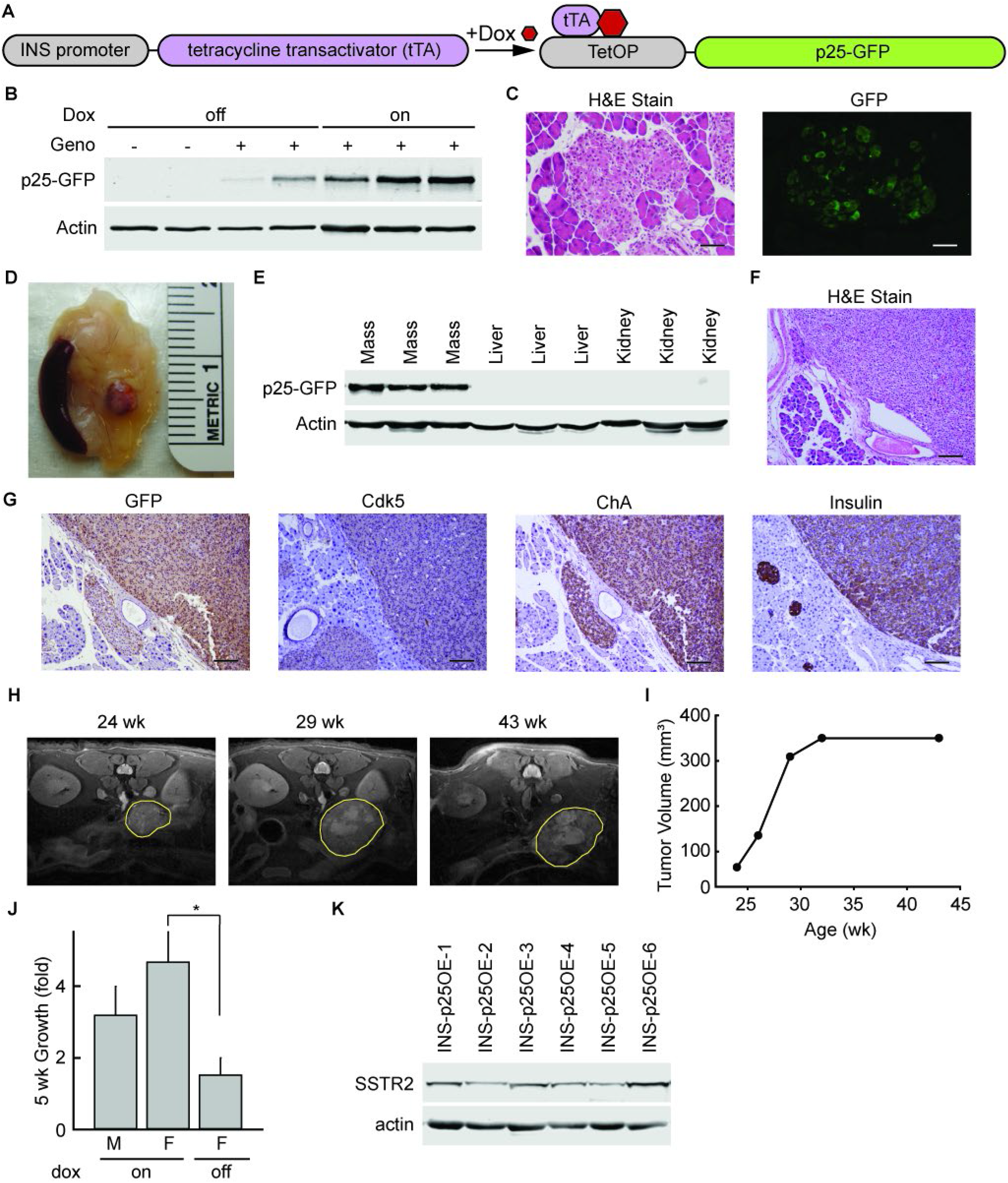
Aberrant activation of Cdk5 generates PanNETs in an inducible bi-transgenic mouse model. A. Schematic of genetic system for regulated tissue-specific expression of p25-GFP. B. Immunoblot for expression of p25GFP in islets isolated from transgene negative (-) and INS-p25OE (+) mice with (on) or without (off) administration of 1 g/L Dox for 4-5 weeks. C. H&E stain and immunofluorescence of sections of pancreas from INS-p25OE mice at 4 weeks post-p25-GFP induction. D. Representative gross image of a pancreatic mass from INS-p25OE mice. E. Immunoblot for expression of p25-GFP in pancreatic mass, liver, and kidney at 12 months induction. F-G. H&E stain (F), and immunostains (G) of primary PanNET from an INS-25OE animal. Scale bars = 100 μm. H. Axial MRI sections from a representative INS-p25OE mouse. PanNET circumscribed in yellow. I. Quantitation of tumor volume over time from a representative INS-p25OE mouse. J. Tumor growth, normalized to initial volume, during the linear growth phase; males (M; n=4) and females (F; n=3 for each group) administered dox since weaning (on) or Dox since weaning followed by discontinuation for 5 weeks at initial tumor detection (off). K. Immunoblot for expression of SSTR2 and actin in p25OE tumors.

Histological analysis of these masses showed a “nesting” pattern in cellular architecture that is characteristic of PanNETs (**Fig. 3F**). Immunoblot and immunostain confirmed the presence of p25GFP and Cdk5 in the lesions (**Fig. 3E, G**). Further, immunostain demonstrated the presence of chromogranin A (ChA), confirming the neuroendocrine phenotype of the lesions. Insulin staining verified the masses were composed of β-cells. In addition, pathological review diagnosed the lesions as well-differientiated PanNETs. These data demonstrate that aberrant activation of the Cdk5 pathway has the potential to directly promote the formation of PanNETs.

To assess growth rate of the INS-p25OE PanNETs, MRI was performed on tumor-bearing mice over a 20-week period beginning when tumors were approximately 50 mm^3^ (**Fig. 3H**). PanNETs in this model exhibited a multiphasic growth pattern. Initial growth was linear with tumors from males and females increasing 3.2-fold and 4.7-fold in size, respectively, over a 5-week timeline (**Fig. 3I-J**). This phase was followed by deceleration and an eventual plateau around 400 mm^3^ (**Fig. 3I**). Removal of dox, to decrease expression of p25GFP after tumor onset, greatly reduced tumor growth rate (**Fig. 3J**).

The presence of a linear growth phase allows detection of changes in tumor growth, in response to experimental therapeutics, in smaller cohorts of animals. To further assess the utility of this model for pre-clinical testing, we examined tumors for the presence of somatostatin receptor 2 (SSTR2), a cell-surface protein commonly overexpressed in human PanNETs and targeted by various FDA-approved treatments for PanNETs. All PanNETs tested exhibited clear SSTR2 expression (**Fig. 3K**).

Human PanNETs present clinically as a highly heterogeneous population of tumors^1,3^. Subgroups of tumors secrete a variety of islet derived hormones while others exhibit no detectable hormone production. To characterize the tumors generated in the new INS-p25OE model, we stained sections of fixed tumors for insulin, glucagon, and somatostatin; three hormones commonly expressed in functional human PanNETs. All PanNETs examined expressed insulin in the tumor mass and a few also exhibited expression of glucagon and somatostatin (**Fig. 4A**).

**Figure 4.**
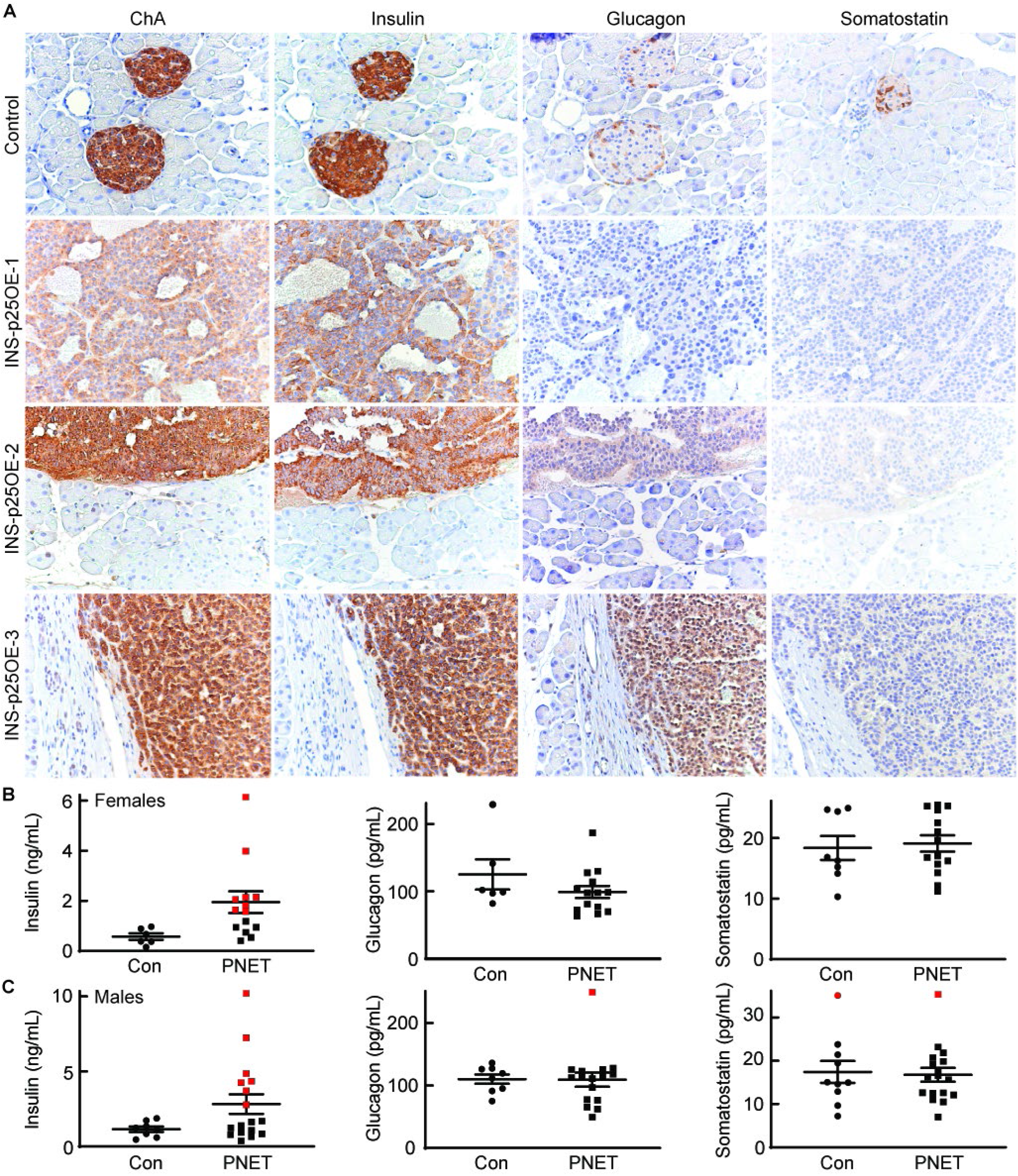
Cdk5 induces both functional and non-functional PanNETs. A. Immunostains of pancreas from control and three representative INS-p25OE animals. B-C. ELISA assays of hormone levels in blood plasma of female (B) and male (C) control (Con; females n=6, males n=8) and tumor-bearing (PNET; females n=14, males n=21) mice. Error bars are SEM; red points illustrate samples with levels that are two SD above the average for controls.

For a tumor to be definitively categorized as clinically functional, in addition to the presence of the hormone in tumor tissue, circulating blood hormones must be elevated to levels capable of inducing physiological effects. Therefore, plasma samples from animals harboring PanNETs and transgene (-) littermates, as controls, were analyzed for insulin, glucagon, and somatostatin. Tumor bearing animals were not found to possess statistically higher average levels of any islet hormone analyzed when assessed collectively (**Fig. 4B-C**).

For higher stringency for classification as non-functional, the data was analyzed again using two standard deviations above the mean of the control group as the cut-off for normal hormones levels. The average insulin levels in normal females and males was statistically different at 0.6 and 1.1 ng/mL, respectively (**Fig. 4B-C**) (p=0.03). Elevations in insulin were present in 57% (8 of 14, red symbols) of tumor-bearing females with 10.8-fold being the highest observed increase relative to control animals. Insulin levels were elevated in 41% (7 of 17, red symbols) of males with 8.9-fold being the highest elevation observed. Normal glucagon levels for females and males were 125 and 110 pg/mL, respectively. Of tumor-bearing animals, only one male exhibited a 2.3-fold elevation of plasma glucagon, less than 1% of the total population and within the natural expected Gaussian distribution. Somatostatin levels in control females and males were 15 and 18 pg/mL, respectively. Both normal and tumor-bearing populations of males contained one animal with somatostatin levels elevated greater than two SD above the mean of the control population, again falling within the natural expected Gaussian curve.

Additionally, we tested the plasma of seven females and seven males lacking large tumor masses but found to possess abnormal islets by histopathological evaluation (data not shown). Insulin was elevated in the plasma of 1 of the 7 additional females. This female also exhibited elevation in somatostatin. One separate female possessed elevated plasma glucagon levels. In males, 2 of the 7 exhibited elevated plasma glucagon, one exhibited elevated plasma insulin, and one exhibited elevated somatostatin. Although immunostaining evaluation identified tumors that were positive for both insulin and glucagon, no animals were found to possess elevation of serum levels of both hormones. One animal, of 45 examined, exhibited elevations in both insulin and somatostatin. Collectively these data demonstrate that 48% of PanNETs generated in the INS-p25OE model are potential insulinomas and 52% do not produce elevations in the serum hormones analyzed and are likely non-functional.

Expression of insulin in all tumors and elevation of circulating insulin levels in 48% of PanNET animals suggested approximately half of the tumors were functional insulinomas. However no pre-mature death was observed in the animals as would be expected from severe hypoglycemia due to overexpression of insulin. To investigate more thoroughly, we tested blood glucose levels in several female and male animals following a 4-6 h fasting window. Surprisingly, only 7% of females (1 of 14) and 23% of males (4 of 17), showed depressed circulating glucose levels under these conditions compared to transgene (-) littermate controls (Supplementary Fig. S2A-B). Because mild insulinemia might take longer to affect glucose levels, we then tested both 4 and 8 h fasting windows in a small set of tumor-bearing females and found that only 17% (1 of 6) exhibited hypoglycemia even after 8 h without food (Supplementary Fig. S2C). Collectively, these data point toward 52-83% of tumors generated from this model being non-functional.

Mutation of the *menin* gene is the most common genetic alteration found in human PanNETs, although the prognostic implications of this mutation are a point of contention. To begin to determine if menin and Cdk5 tumorigenic pathways overlap, we analyzed the presence of menin, Cdk5, p35, and downstream components of the menin pathway in PanNETs from the MEN^+/-^ model (MEN) and the INS-p25OE model (**Fig. 5A**). As expected, levels of menin were reduced in MEN^+/-^ tumors. Analysis of the downstream targets of menin, p18^Ink4c^ and p27^KIP1^, also revealed decreased expression in MEN tumors compared to INS-p25OE tumors. This comparison suggests that aberrant activation of the Cdk5 pathway does not lead to inhibition of genes targeted by menin.

**Figure 5.**
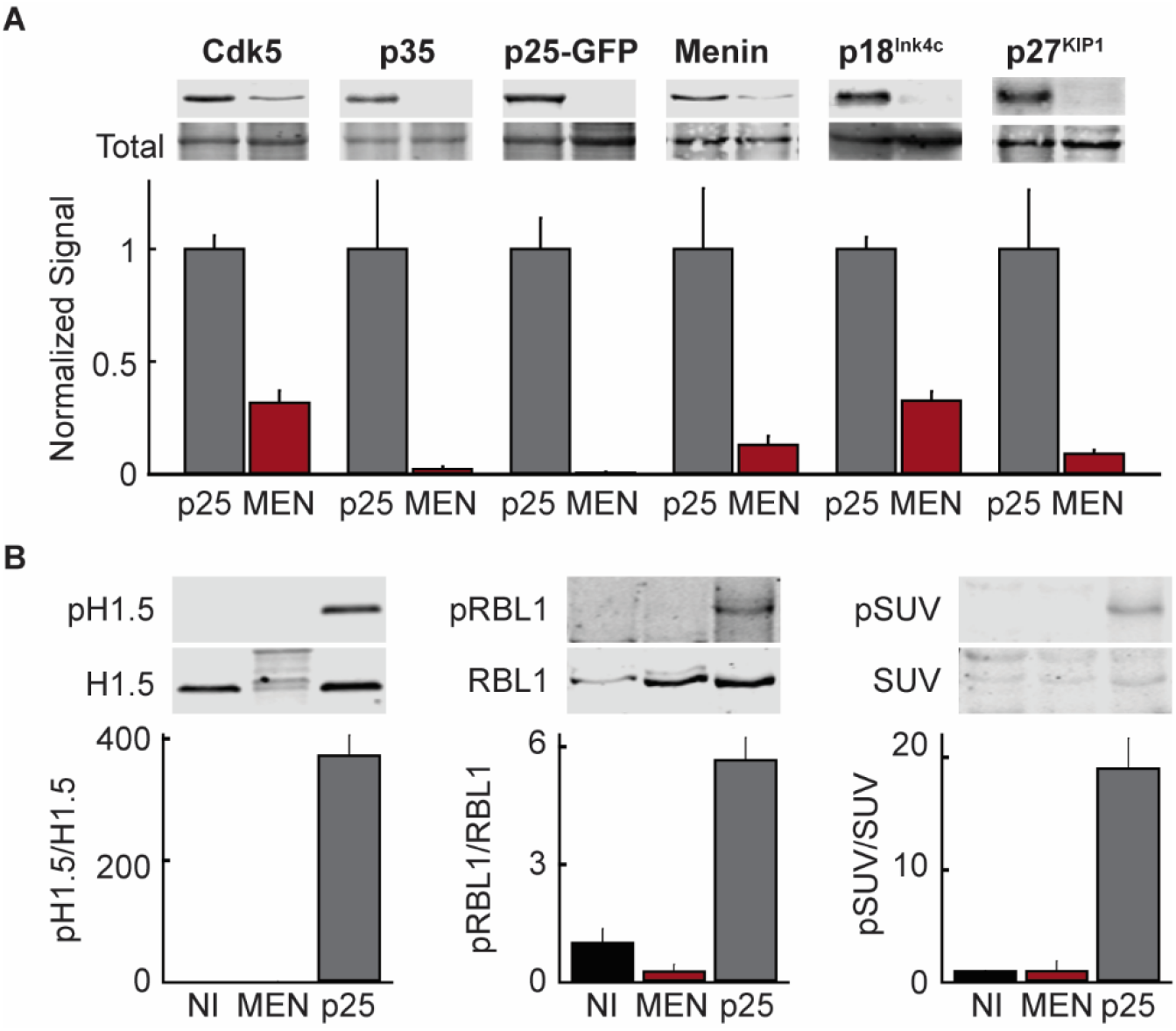
Cdk5 and menin pathways are distinct drivers of mPanNETs. A. Quantitative immunoblot of Cdk5 pathway components, menin, and downstream targets of menin in MEN^-/+^ tumors (MEN; n=4) and INS-p25OE tumors (p25; n=5). B. Quantitative immunoblot of downstream targets of Cdk5 in normal mouse islets (NI; n=3), MEN^-/+^ tumors (MEN; n=4), and INS-p25OE tumors (p25; n=7); phosphorylated Ser18-H1.5 (pH1.5), phosphorylated Ser988-RBL1 (pRBL1), phosphorylated Ser391-SUV39H1 (pSUV).

Levels of Cdk5 and p35 were also reduced in MEN^+/-^ tumors, suggesting that PanNETs arising from loss of function mutations in *menin* are not driven by aberrant activation of Cdk5. To explore this observation further, we interrogated phosphorylation levels of three proteins previously identified as downstream targets of aberrant Cdk5 in thyroid neuroendocrine tumors: phospho-Ser18 histone H1.5, Ser988 RBL1, and Ser391 SUV3H1^17^. Interestingly, each of these markers was highly phosphorylated in INS-p25OE tumors. In contrast, these signals were almost completely absent in normal islets as well as MEN^+/-^ tumors, further supporting that loss of menin does not lead to aberrant activation of Cdk5 as a part of its tumorigenic process (**Fig. 5B**). Together, these data indicate that menin and Cdk5 pathways constitute separate and independent tumorigenic pathways.

While these studies show that tumors retain dependence upon Cdk5 activity for sustained growth, the variability in age of onset combined with 75% penetrance by 12 months of age (Supplementary Fig. S1) raises the possibility that additional alterations occur and facilitate tumor formation. To investigate this further, we performed whole exome sequencing on five INS-p25OE PanNETs; three functional and two non-functional tumors. Interestingly, high heterogeneity was observed in the genetic landscape of the these tumors as is also found in human tumors (**Fig. 6**). Several classes of mutations were observed throughout multiple chromosomes including alterations in introns, exons, 3’ UTRs, and 5’ UTRs (**Fig. 6A**). Single nucleotide polymorphisms (SNPs) were the most common type of alteration detected (**Fig. 6B**). Examination of mutations from translated regions revealed very little overlap among samples (**Fig. 6C**). Although mutations in identical genes among INS-p25OE tumors were rare, alterations in genes encoding regulatory subunits of the PIK3 pathway were found in three of the five samples. Mutations in the catalytic subunit of PIK3 are known to be enriched in human PanNETs^13^. This finding prompted a full comparison with sequencing datasets from human PanNETs, which revealed that 48 genes with mutations in INS-p25OE tumors are also mutated in a published set of 98 human PanNETs^40^ (**Fig. 6D** and Supplemental Table S2). Together, the analyses indicate that the INS-p25OE model shares appreciable genetic overlap with human PanNETs.

**Figure 6.**
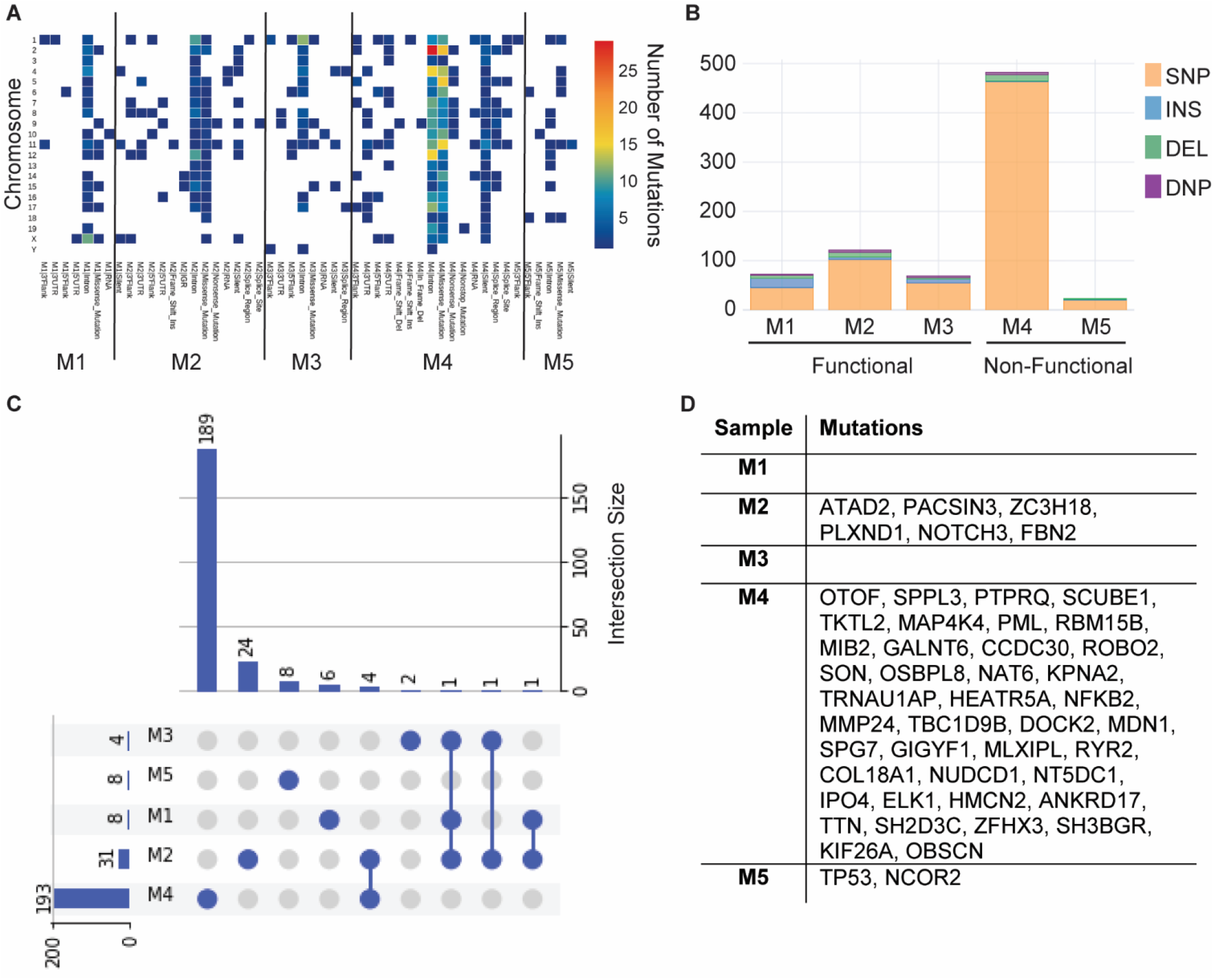
INS-p25OE tumors possess mutations found in human PanNETs. A. Heat map of number of total mutations in each INS-p25OE PanNET by classification per chromosome. B. Type of mutations present in each INS-p25OE PanNET. C. Waterfall plot of overlapping exonal mutations among INS-p25OE PanNETs. D. Table of mutations in INS-p25OE PanNETs found to overlap with mutations in human PanNETs.

To further understand the molecular changes that lead to tumor development in the INS-p25OE model, we performed mRNA sequencing on six INS-p25OE PanNETs, three functional and three non-functional tumors, and compared levels of gene expression to that observed in normal mouse islets (**Fig. 7A**). Interestingly, higher heterogeneity was observed in the non-functional group than in the functional group (**Fig. 7B**). Comparing the total tumor group to normal islets, we found that 796 genes were upregulated while 533 genes were downregulated (**Fig. 7C**). Of note, genes such as BRCA2, STAT4 and TOP2A were dysregulated, similar to previous observations from human PanNETs (**Fig. 7D**)^41,42^. Ingenuity Pathway Analysis revealed upregulation of four pathways that relate to cell cycle regulation, one pathway that involves DNA repair, one that is important for vascularization, and three that are linked to collagen and extracellular matrix regulation (**Fig. 7E**).

**Figure 7.**
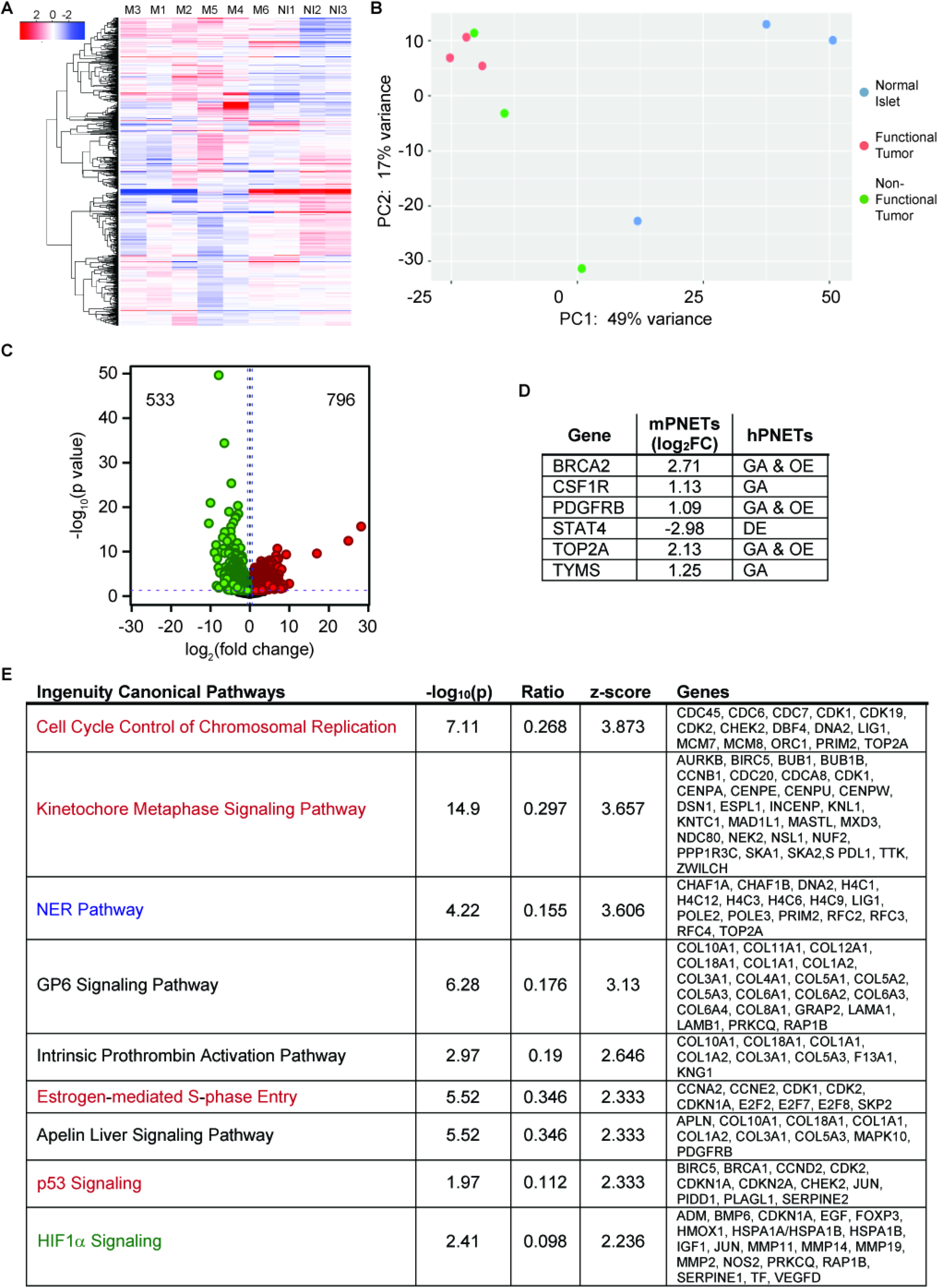
RNA-seq. A. Heatmap of differentially expressed genes in normal mouse islets (NI1-3), functional (M1-3) and non-functional (M4-6) INS-p25OE PanNETs. B. Principal component analysis of expression data. C. Volcano plot of annotated genes upregulated and downregulated compared to normal mouse islets. D. Table of differentially expressed genes that overlap with alterations in human PanNETs; gene amplification (GA), overexpression (OE), decreased expression (DE). E. Ingenuity Pathway Analysis of differentially expressed genes. Pathways are related to cell cycle (red), DNA repair (blue), vascularization (green), and extracellular matrix (black).

Although the INS-p25OE model generates genetically (**Fig. 6**) and phenotypically (**Fig. 4**) heterogenous tumors as is observed in human patients, heterogenous models require large cohort sizes to identify responses in pre-clinical trials. In addition, the primary model requires 6-12 months to form tumors. Therefore, we established tumor-dervied allografts from INS-p25OE primary PanNETs as second tool that could be utilized for quick screening in a large, homogenous cohort of animals. We implanted 2 mm x 2 mm sections of tissue from a primary tumor (P0) into five recipient BL/6 male mice. Allograft tissue established new tumors (P1) with 100% penetrance and, on average, within 17 weeks, reducing the timeframe for development from 45 weeks in P0 mice to 17 weeks in P1 animals (**Fig. 8A-D**). Further, allografts can be serially passaged with 100% penetrance and establish 3^rd^ generation tumors (P2), on average, within 8 weeks (**Fig. 8C-D**). Allografts retain expression of the p25-GFP transgene and tumors grow 4.3 fold in a 5 week period, very similar to growth rates of primary PanNETs (**Fig. 8B and E, Fig. 3J**). Allografts retain the well-differientiated neuroendocrine phenotype of the primary tumors, including tumor architecture and positivity for ChA and insulin staining. (**Fig. 8F**).

**Figure 8.**
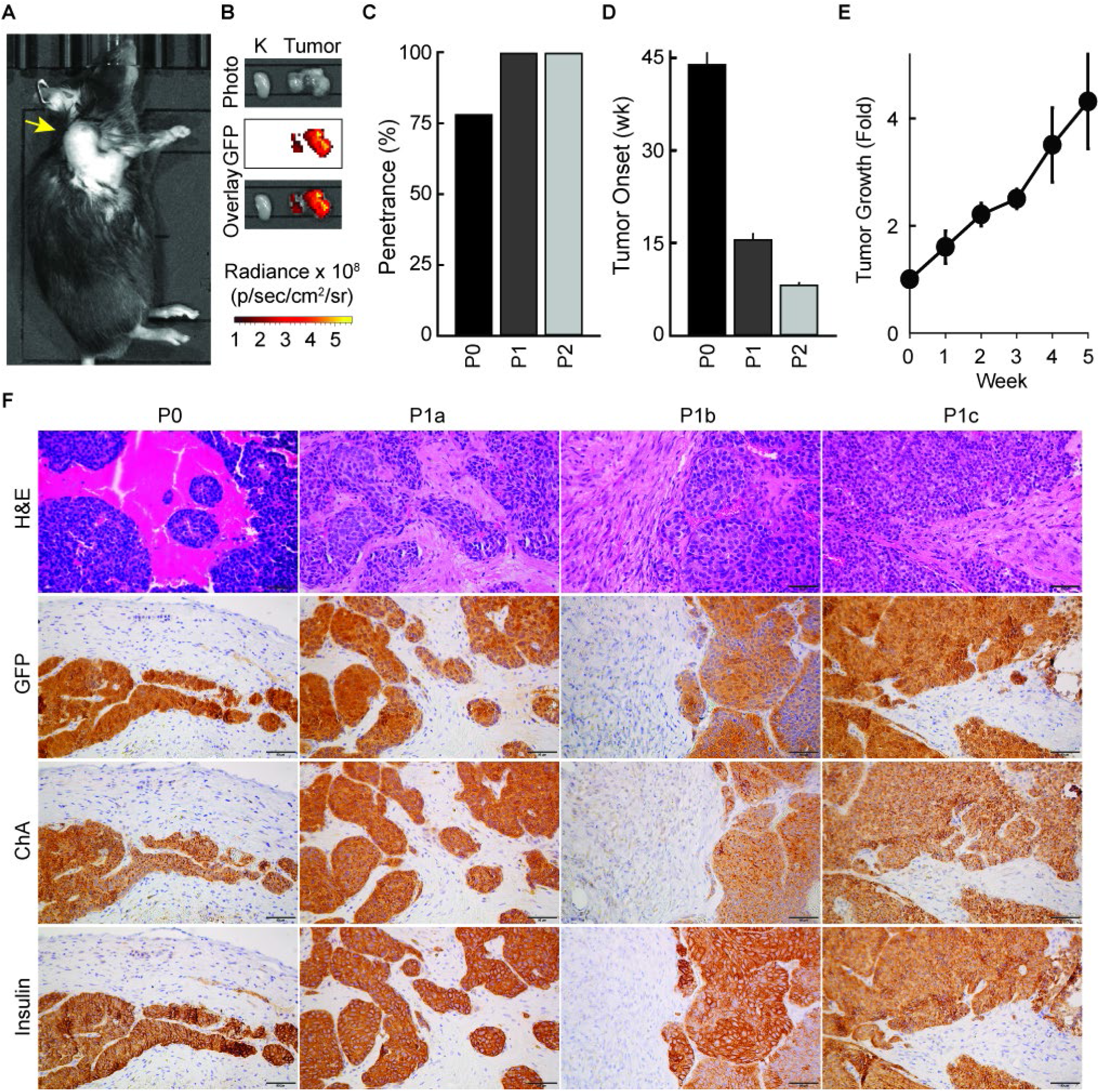
INS-p25OE PanNETs form successful allografts. A.Photograph from IVIS imagine of representative allograft tumor model; tumor marked with yellow arrow. B. Photograph and fluorescence imaging from IVIS, ex vivo, of kidney (left) and tumor (right) from representative allograft model. C. Penetrance in primary INS-p25OE model (P0), passage 1 models (P1), and passage 2 models (P2). D. Tumor onset in the same groups as C. E. Fold growth of passage 1 tumors over a 5 week period starting at approximately 100 mm^3^ (week 0). F. H&E stains and immunostains of primary INS-p25OE model (P0) and three passage 1 models (P1a-c).

## Discussion

Progress in the development of therapeutics that specifically target NETs has been hampered in part by an insufficient number of animal models in which to perform preclinical experimentation. While PanNETs co-occur with multiple other tumor types in diverse genetically engineered mouse models, only two main types of transgenic mouse models have been generated and utilized for pre-clinical PanNET studies prior to the development of the INS-p25OE model reported here^43,44^. The MEN^+/-^ conventional knockout model develops PanNETs, as well as parathyroid and pituitary NETs, and has been utilized to explore new therapeutics such as anti-VEGF-A monoclonal antibody therapy and Pasireotide for efficacy toward PanNETs^43,45,46^. This model is expected to be especially relevant to the approximately 40% of PanNET patients that possess a mutation in the gene menin. Both pan-pancreas and islet-specific conditional homozygous knockouts of the menin gene also produce PanNETs^43^. Of note, all of the PanNETs from these menin knockout models are insulinomas or gastrinomas while approximately 85% of human PanNETs are classified as non-functional. Therefore, additional models would be highly beneficial.

A second conditional transgenic mouse model of PanNETs is the RIP1-Tag2 line^47^. This model was generated by cloning the large T-antigen of SV40, a known oncogenic driver, downstream of the rat insulin promoter for expression in β-islet cells. This model develops aggressive insulinomas, including both well- and poorly-differentiated subsets, and has been successfully utilized to explore new therapeutics such as sunitinib and mTOR inhibitors^48-50^. Interestingly, crossing the RIP1-Tag2 mouse model into the A/J background leads to formation of tumors that do not express insulin^51^. The A/J background has a known SNP, relative to the C57BL/6 background, in the *Insm1* gene. *Insm1*, which encodes a transcription factor that promotes neuroendocrine differentiation and is required for insulin expression in β cells, was implicated in the loss of insulin expression observed in the model^52^. Development of this model will undoubtedly provide insight into non-functional tumor physiology. However, these tumors are more poorly differentiated than tumors from the parent C57BL/6 background and the population of human tumors to which it is relevant will need to be carefully investigated as rare, poorly-differentiated G3 neuroendocrine carcinomas, and relatively more abundant, well-differentiated G3 NETs, are molecularly distinct tumor types^51,53^.

Here, we present development of a novel, dox-inducible, conditional mouse model of PanNETs in which activation of the Cdk5 pathway in β-islet cells leads to slow growing islet tumors with heterogeneous hormone production profiles, including a large subset of non-functioning, well-differentiated tumors. The utility of this model is further extended by the ability to generate multiple allograft animals from each primary PanNET. As these second generation animals also possess a fully functional immune system, this method for generating large homogenous cohorts of immunocompetent PanNET models will be especially useful for exploration of immunotherapies, a modality whose implementation has lagged for neuroendocrine cancers.

Male and female cohorts were interrogated as separate groups when characterizing the INS-p25OE primary PanNET model so that differences linked to sex could be uncovered. Surprisingly, although females exhibited a higher propensity for elevation of plasma insulin levels compared to males, fewer females developed hypoglycemia in response to fasting. This may be due to the fact that total insulin levels were higher in “elevated” males than “elevated” females. We have found no clinical analysis of human populations that indicate that non-functioning PanNETs are more common in one sex versus the other, although NETs in general are slightly more common in females^4^.

The INS-p25OE model reported here is molecularly distinct from the MEN^+/-^ model and likely represents a group of human PanNETs in which mutation of the gene menin is not the key driving factor. Although causative events that lead to Cdk5 pathway activation in humans are unclear, Cdk5 misregulation has been demonstrated in multiple types of human NETs^17,18^. In addition, genetic and transcriptional data point to multiple overlaps between human tumors and the INS-p25OE model. Significant overlap was also observed at the functional level, as ∼85% of human tumors are non-functional and we observed a similar distribution of functional and non-functional tumors in the INS-p25OE model. This newly developed model will serve as a useful platform for molecular characterization of the population of human PanNETs in which aberrant activation of Cdk5 is present as well as the development and testing of new therapeutics that target those pathways. Moreoever, because this model more faithfully reflects human PanNET biology, it will facilitate development of a variety of therapeutic strategies, not limited to targeting of Cdk5.

## Methods

### Human tissue collection

Samples were collected in accordance with institutional review board (IRB) regulations under Louisiana State University IRB 5774 and University of Alabama at Birmingham IRB 300002147.

### Histology

Tissues were fixed in formalin, embedded in paraffin, and sliced into 5 μm sections for placement on glass slides. Samples were deparaffinized and subjected to high temperature antigen retrieval in citrate buffer (pH 6.0). For immunostaining, samples were permeabilized in 0.3% Triton X-100, incubated in 0.3% hydrogen peroxide, blocked with 3% normal goat serum, and then incubated overnight at 4°C in primary antibodies. Human and mouse tissue was immunostained for Cdk5 (PhosphoSolutions 308-Cdk5; 1:50) and p35/p25 (Santa Cruz sc-820; 1:50). Mouse tissue was immunostained for GFP (Cell Signaling Technology 2956; 1:200), ChA (Abcam ab15160; 1:500), insulin (Abcam ab63820; 1:2000), glucagon (Santa Cruz sc7779; 1:200), somatostatin (Abcam ab108456; 1:450). Biotinylated secondary antibodies (Pierce 31820 or 31800; 1:500) were applied to slides for 1 h at room temperature followed by 30 min of HRP streptavidin. Slides were then incubated with DAB Chromogen (Dako Liquid DAB+ substrate K3468) and counter stained with hematoxylin. Standard procedures were used for H&E staining. The human PanNET TMA was prepared by the UAB Research Pathology Core. Slides were immunostained as stated above. Images were deconvoluted using Fiji ImageJ. The mean intensity of a fixed region of interest for each core in the resulting DAB channel was measured and then converted to optical density using the formula: OD = Log (Max intensity/mean intensity) for semi-quantitative analysis.

### Cell Culture

All cells were cultured in a humidified incubator at 37°C under 5% CO_2_. Fibroblasts were grown in DMEM plus 10% FBS. BON and QGP cells were grown in RPMI plus 10% FBS, 100 μg/ml penicillin, and 100 μg/ml streptomycin. NT3 and NT18 cells were cultured in RPMI 1640 GlutMAX plus 10% FCS, 20 ng/ml EGF, 10 ng/ml FGF2, 100 μg/ml penicillin, and 100 μg/ml streptomycin.

### Cell growth assay

Cells were seeded onto 96-well plates and allowed to adhere for 24 h. Cells were then treated twice (day 1 and day 3) with various concentrations of inhibitor, as shown, and viability measured after 5 days by MTT assay. IC_50_ values were determined by 4-parameter logistic regression.

### INS-p25OE animal model

All animal work was performed in accordance with the Animal Welfare Act and the Guide for the Care and Use of Laboratory Animals under UTSW and UAB Institutional Animal Care and Use Committee approved protocols. Bi-transgenic INS-p25OE animals were generated from crossing of the tetOp-p25GFP strain (The Jackson Laboratory stock # 005706) with the Ins2-rtTA strain (Provided by Dr. Alvin C. Powers at Vanderbilt; available from The Jackson Laboratory stock # 008250). Breeders and pups were maintained in the absence of doxycycline to allow for normal development of offspring prior to transgene induction. Upon weaning, at 3-4 weeks of age, offspring were administered 1 mg/L doxycycline via drinking water to induce transgene expression in bi-transgenic animals. Bi-transgenic animals were co-housed with transgene negative littermates. Transgene negative littermates were used as normal controls. All mice were maintained in the C57BL/6 background. Animals were euthanized by CO_2_ administration and cardiac perfusion.

### MRI

MRI was performed with a Bruker Biospec 9.4 Tesla instrument using Paravision 5.1 software (Bruker Biospin, Billerica, MA). A Bruker 72 mm ID volume coil was used for excitation and a custom 24 mm surface coil for signal reception (Doty Scientific Inc., Columbia, SC). Mice were anesthetized with isoflurane gas and respiration observed with a MRI-compatible physiological monitoring system (SA Instruments Inc., Stony Brook, NY). Animals were imaged in supine position on a Bruker animal bed system with circulating heated water to maintain body temperature. A 2D T2-weighted RARE sequence was used for imaging of the abdomen. The following imaging parameters were used: TR/TE = 2000/25 ms, echo spacing = 12.5 ms, ETL = 4, 2 averages, 29 contiguous axial slices with 1 mm thickness, FOV = 30×30 mm and matrix = 300×300 for an in-plane resolution of 100 μm. Prospective respiratory gating was used to minimize motion artifacts. Tumors volumes were quantitated using ImageJ software.

### Immunoblot

Cells were lysed in 1% SDS plus 50 mM NaF. Samples were sonicated briefly, spun at 20,000 g for 5 min, and supernatant combined with Laemmli buffer for analysis by SDS-PAGE followed by transfer to PVDF for immunoblotting. Tumors were crushed while frozen and then processed using the same protocol. Immunoblotting was performed using antibodies for Cdk5 (Rockland 200-301-163; 1:1000), p35 (Santa Cruz sc-820; 1:300), GFP (Cell Signaling Technology 2956; 1:2000), SSTR2 (Santa Cruz sc-365502; 1:500), Menin (Santa Cruz sc-374371; 1:250), p18Ink4c (Invitrogen 393400; 1:500), and p27Kip1 (Cell Signaling Technology 2552; 1:1000), pS18H1.5 (Bibb Lab; 1:1000), H1.5 (Santa Cruz sc-247158; 1:1000), pS988RBL1 (Bibb Lab; 1:1000), RBL1 (Santa Cruz sc-318; 1:500), pS392-SUV39H1 (Bibb Lab; 1:300), SUV39H1 (Sigma S8316; 1:500), and actin (Invitrogen AM4302; 1:5000). Revert 700 Total Protein Stain (LICOR 926-11011) was used per manufacturer’s protocol.

### Whole Exome Sequencing

The analysis of raw WES data was performed using MoCaSeq pipeline (source code: https://github.com/roland-rad-lab/MoCaSeq). The pipeline was set up using the docker container and Ubuntu Linux. Specifically, the raw reads were trimmed aligned to the mouse reference genome GRCm38.p6 using Trimmomatic 0.38 and BWA-MEM 0.7.17, respectively. For further post-processing, Picard 2.20.0 and GATK 4.1.0.0 were used. For the loss of heterozygosity (LOH) analyses from WES data, somatic SNP calling was performed using Mutect2. To avoid ambiguous SNP positions resulting from mis-mapping, only reads with a mapping quality of 60 were kept in LOH analyses. For CNV calling, CopywriteR 2.6.1.216 was used, which extracts DNA copy number information from targeted sequencing by utilizing off-target reads. Finally, the downstream analysis and visualization were done using custom Python (v.3.8) and Shell scripting. Data from mice were compared to human data deposited with the European Genome-Phenome Archive under EGAD00001002684.

### RNASeq Analysis

RNA was isolated from tissue using RNeasy Plus Mini Kit (Qiagen 74134). RNA was transcribed to cDNA using NEBNext Ultra™ RNA Library Prep Kit for Illumina (NEB E7530). RNA sequencing was performed using single-end 75 bp reads on an Illumina NextSeq500. The RAW sequences were trimmed using Trimmomatic 0.38 and low-quality reads were removed. The quantification of the expression of transcripts of preprocessed sequences was using salmon 1.4.0 and mm10 mouse reference genome. The resulting quant (transcript abundance estimates) values were utilized for the differential expression analysis. Differential gene expression analysis was done using DESeq2 and for downstream analysis and visualization python (v.3.8) and Bash scripting were used.

### Allograft models

Primary tumors were removed from INS-p25OE mice and diced into ∼2 mm x 2 mm sections. These sections were implanted into both the right and left flanks of C57BL/6 P1 (passage 1) recipient mice by trocar. Tumor size was monitored by measurement with calipers. P2 mice were generated by passaging P1 tumors into a second generation of C57BL/6 recipient mice.

### Statistical Analysis

Comparisons between two groups were performed using two-tailed Student’s *t*-test. Comparisons between three groups were performed using one-way ANOVA. Sample sizes are provided within figure legends or in results. (*p<0.05).

## Acknowledgments

Ins2-rtTA mice were kindly provided by Dr. Alvin C. Powers (Vanderbilt University). PanNETs from the MEN^+/-^ mouse model (18-22 months old mice) were kindly provided by Vaishali Parekh of Dr. Sunita K. Agarwal’s lab (NIH/NIDDK). We thank the Pathology Core at UAB for TMA production, the UAB Small Animal Imaging Facility for MRI on mice, and the Heflin Center for Genomic Science at UAB for WES and RNAseq. We thank Boehringer-Ingelheim and Frank Gillardon for providing Indo A. This research was further supported by core capabilities provided by the O’Neal Comprehensive Cancer Center.

## Author Contributions

A.M.C. and J.A.B. conceptualized the study. A.M.C., B.H., C.T., R.G., and W.H. performed immunostaining. A.M.C. performed biochemistry, immunoblots, ELISA assays, analysis of MRIs, quantitation of immunostaining, and data interpretation. F.V. performed cell growth assays. R.T. maintained the mouse colony, assisted with tissue harvesting and molecular biology. A.M.C., T.M., C.G., and J.B.R. generated allografts. H.B. harvested islets from mice. C.T. and E.K. performed pathological assessment of tumors. J.T. developed the MRI protocol. N.K. and M.S.M. performed bioinformatics analyses. A.M.C. and J.A.B. assembled figures and wrote the manuscript. R.B., H.C., J.S., J.B.R., M.S.M., and J.A.B. supervised the study. All authors reviewed and edited the manuscript.

