## Supplemental Data for "Cdk5 drives formation of heterogeneous pancreatic neuroendocrine tumors"

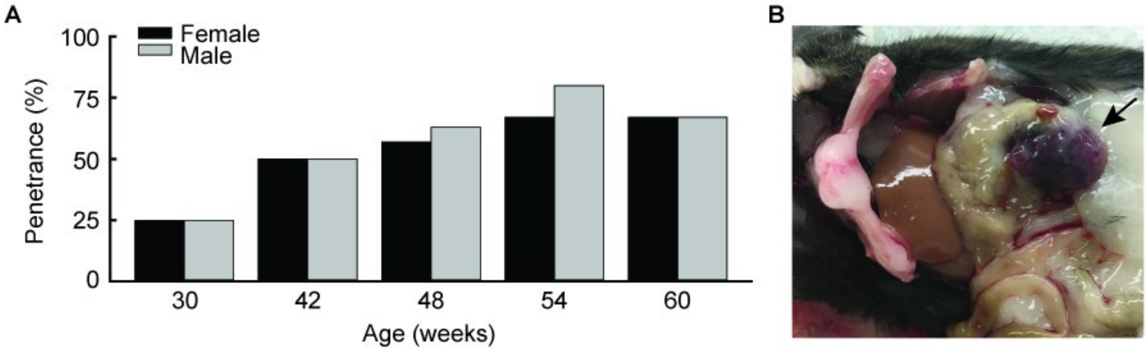


**Supplemental Figure S1. Penetrance of PanNETs in INS-p25OE model.** A. Detection of visible tumors by autopsy (n=4-8). B. Representative image of PanNET in autopsy; tumor marked with black arrow.

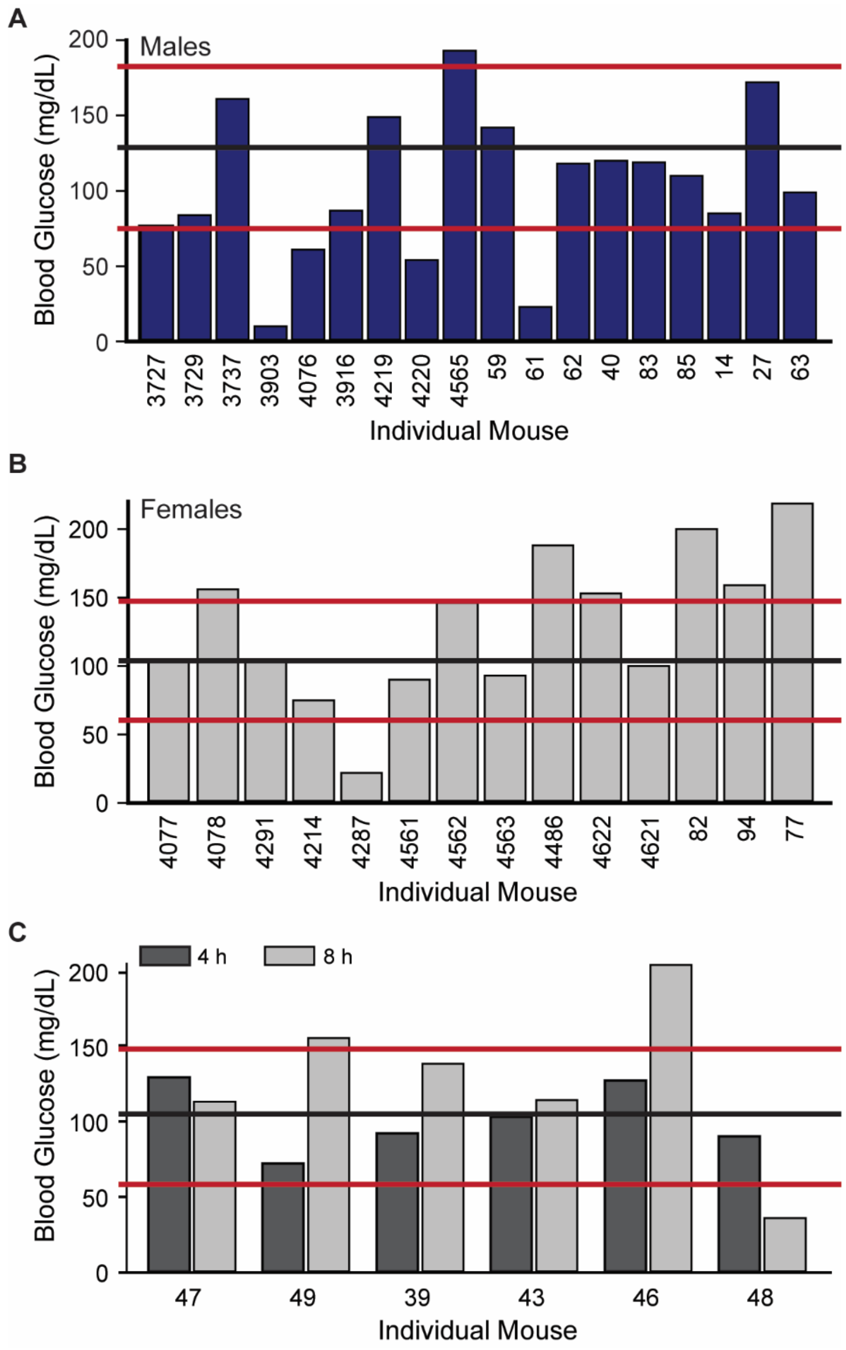


**Supplemental Figure S2. INS-p25OE animals exhibit low penetrance of hypoglycemia.** A-B. Blood glucose levels in male (A) and female (B) INS-p25OE tumor-bearing animals (n=18 and 14, respectively) after 4 h fast. C. Blood glucose levels in female INS-p25OE tumor-bearing animals after 4 and 8 h fast (n=6). Black lines represent average glucose levels in control littermates. Red lines denote two standard deviations from average of controls.

**Supplemental Table S1. Map of human PanNET TMA.**

| | | | | | | | | |
|---|---|---|---|---|---|---|---|---|
| Normal<br>Spleen | WD-NET<br>NF<br>Grade 1 | WD-NET<br>NF<br>Grade 1 | WD-NET<br>NF<br>Grade 1 | WD-NET<br>NF<br>Grade 2 | WD-NET<br>NF<br>Grade 1 | WD-NET<br>NF<br>Grade 1 | WD-NET<br>NF<br>Grade 2 | WD-NET<br>NF<br>Grade 1 |
| Normal<br>Liver | WD-NET<br>NF<br>Grade 1 | WD-NET<br>NF<br>Grade 2 | WD-NET<br>NF<br>Grade 2 | WD-NET<br>NF<br>Grade 2 | WD-NET<br>NF<br>Grade 2 | WD-NET<br>NF<br>Grade 2 | WD-NET<br>NF<br>Grade 1 | WD-NET<br>NF<br>Grade 1 |
| Normal<br>Prostate | WD-NET<br>NF<br>Grade 2 | WD-NET<br>NF<br>Grade 1 | WD-NET<br>NF<br>Grade 1 | WD-NET<br>NF<br>Grade 1 | WD-NET<br>NF<br>Grade 2 | WD-NET<br>NF<br>Grade 1 | WD-NET<br>NF<br>Grade 1 | Blank |
| Normal<br>Placenta | WD-NET<br>NF<br>Grade 1 | WD-NET<br>NF<br>Grade 1 | WD-NET<br>NF<br>Grade 2 | WD-NET<br>NF<br>Grade 1 | WD-NET<br>NF<br>Grade 1 | WD-NET<br>NF<br>Grade 1 | WD-NET<br>Insulinoma<br>Grade 2 | WD-NET<br>Insulinoma<br>Grade 1 |
| Normal<br>Tonsil | WD-NET<br>NF<br>Grade 3 | Blank | WD-NET<br>NF<br>Grade 2 | WD-NET<br>NF<br>Grade 1 | WD-NET<br>NF<br>Grade 2 | WD-NET<br>NF<br>Grade 1 | WD-NET<br>NF<br>Grade 1 | Blank |

*WD - Well-differentiated, NF – Non-functional

**Supplemental Table S2. Mutation frequency, in humans, for genes mutated in INS-p25OE PanNETs.**

| SYMBOL | GENE_ID | Frequency (%) in Humans |
|---|---|---|
| TP53 | ENSG00000141510 | 4 |
| TTN | ENSG00000155657 | 5 |
| OBSCN | ENSG00000154358 | 3 |
| FBN2 | ENSG00000138829 | 3 |
| SH3BGR | ENSG00000185437 | 2 |
| COL18A1 | ENSG00000182871 | 2 |
| NUDCD1 | ENSG00000120526 | 2 |
| RYR2 | ENSG00000198626 | 4 |
| NAT6 | ENSG00000243477 | 2 |
| PTPRQ | ENSG00000139304 | 2 |
| MMP24 | ENSG00000125966 | 2 |
| NOTCH3 | ENSG00000074181 | 2 |
| NT5DC1 | ENSG00000178425 | 2 |
| OSBPL8 | ENSG00000091039 | 1 |
| ATAD2 | ENSG00000156802 | 1 |
| HEATR5A | ENSG00000129493 | 1 |
| MLXIPL | ENSG00000009950 | 1 |
| SH2D3C | ENSG00000095370 | 1 |
| RBM15B | ENSG00000179837 | 1 |
| TBC1D9B | ENSG00000197226 | 1 |
| NFKB2 | ENSG00000077150 | 1 |
| SON | ENSG00000159140 | 1 |
| MAP4K4 | ENSG00000071054 | 1 |
| SPPL3 | ENSG00000157837 | 1 |
| PML | ENSG00000140464 | 1 |
| ANKRD17 | ENSG00000132466 | 1 |
| TKTL2 | ENSG00000151005 | 1 |
| KPNA2 | ENSG00000182481 | 1 |
| ROBO2 | ENSG00000185008 | 1 |
| GIGYF1 | ENSG00000146830 | 1 |
| ELK1 | ENSG00000126767 | 1 |
| IPO4 | ENSG00000196497 | 1 |
| OTOF | ENSG00000115155 | 1 |
| ZC3H18 | ENSG00000158545 | 1 |
| KIF26A | ENSG00000066735 | 1 |
| ZFHX3 | ENSG00000140836 | 1 |
| SPG7 | ENSG00000197912 | 1 |
| PLXND1 | ENSG00000004399 | 1 |
| SCUBE1 | ENSG00000159307 | 1 |
| MIB2 | ENSG00000197530 | 1 |
| HMCN2 | ENSG00000148357 | 1 |
| NCOR2 | ENSG00000196498 | 1 |
| TRNAU1AP | ENSG00000180098 | 1 |
| GALNT6 | ENSG00000139629 | 1 |
| MDN1 | ENSG00000112159 | 1 |
| PACSIN3 | ENSG00000165912 | 1 |
| CCDC30 | ENSG00000186409 | 1 |
| DOCK2 | ENSG00000134516 | 1 |
